# Safety and Efficacy of Avaren-Fc Lectibody Targeting HCV High-Mannose Glycans in a Human Liver Chimeric Mouse Model

**DOI:** 10.1101/2020.04.22.056754

**Authors:** Matthew Dent, Krystal Hamorsky, Thibaut Vausselin, Jean Dubuisson, Yoshinari Miyata, Yoshio Morikawa, Nobuyuki Matoba

## Abstract

Infection with hepatitis C virus (HCV) remains to be a major cause of morbidity and mortality worldwide despite the recent advent of highly effective direct-acting antivirals. The envelope glycoproteins of HCV are heavily glycosylated with a high proportion of high-mannose glycans (HMGs), which serve as a shield against neutralizing antibodies and assist in the interaction with cell-entry receptors. However, currently there is no approved therapeutic targeting this potentially druggable biomarker. Here, we investigated the therapeutic potential of the lectibody Avaren-Fc (AvFc), a HMG-binding lectin-Fc fusion protein. *In vitro* assays showed AvFc’s capacity to neutralize cell culture-derived HCV in a genotype independent manner with IC_50_ values in the low nanomolar range. A histidine buffer-based AvFc formulation was developed for in vivo studies using the PXB human liver chimeric mouse model. Systemic administration of AvFc was well tolerated; after 11 consecutive doses every other day at 25 mg/kg, there were no significant changes in body or liver weights, nor any impact noted in blood human albumin levels or serum alanine aminotransferase activity. Gross necropsy and liver pathology further confirmed the lack of discernible toxicity. This treatment regimen successfully prevented genotype 1a HCV infection in all animals, while an AvFc mutant lacking HMG binding activity failed to block the infection. These results suggest that targeting envelope HMGs is a promising therapeutic approach against HCV infection. In particular, AvFc may provide a safe and efficacious means to prevent recurrent infection upon liver transplantation in HCV-related end-stage liver disease patients.

## INTRODUCTION

Hepatitis C virus (HCV) is an enveloped monopartite positive sense ssRNA virus in the family *Flaviviridae* and the causative agent of hepatitis C disease. Its genome encodes three structural (core, E1, E2) and seven non-structural proteins (p7, NS2, NS3, NS4A, NS4B, NS5A, NS5B) (1). HCV is highly heterogenous and globally distributed, consisting of seven genotypes each further subdivided into multiple subtypes. Genotype 1 is the predominant genotype worldwide and particularly concentrated in high-income and upper-middle income countries, whereas genotype 3 and 4 are more common in lower-middle and low-income countries (2). In the United States, injection drug use represents the primary risk factor for contracting HCV infection (3, 4). Around 15-25% of people acutely infected with HCV will clear the virus, while the remainder will develop chronic infection that can persist largely unnoticed for decades. Indeed, many HCV carriers discover their chronic infection after they have developed cirrhosis (5). Chronic HCV infection is also associated with the development of hepatocellular carcinoma, and those with the disease are more likely to develop cryoglobulinemia and non-Hodgkin’s lymphoma (6).

There is no vaccine currently available for HCV. Prior to 2011, the standard chronic HCV treatment was a non-specific antiviral medication using ribavirin combined with a pegylated interferon-α, which was associated with significant toxicity and limited treatment efficacy (7). In 2011, the U.S. Food and Drug Administration approved the first generation of direct-acting antivirals (DAAs) for HCV: boceprevir and telaprevir, both of which inhibit the viral protease (NS3/4A) but required cotreatment with ribavirin and peginterferon (8, 9). Further approval of more potent DAAs, such as NS3/4A, NS5B and NS5A inhibitors led to the development of oral ribavirin/peginterferon-free regimens (5). Multi-DAA regimens achieve sustained virologic response (defined as a period of time with no viral RNA detection) rates as high as 100%, and are less toxic and more tolerable than their predecessors (10-13). While the cure rates are remarkable, there exist populations of patients who may not benefit from DAA therapy (14), especially patients with decompensated cirrhosis due to chronic HCV infection, for whom liver transplantation may be a last resort (15). Moreover, recurrent infection occurs universally and rapidly post liver transplantation (16, 17), which increases the risk of accelerated cirrhosis, graft failure and death (18). DAAs, by their nature, cannot prevent recurrent infection. Therefore, alternative or complementary therapies to DAAs that can block viral entry to target cells, such as antibodies or other molecules acting alike, may need to be considered in these circumstances (18, 19). However, there is currently no entry inhibitor approved for HCV treatment.

The HCV envelope proteins E1 and E2 are heavily glycosylated and, like glycoproteins of other enveloped viruses (HIV and the coronaviruses, for instance), have a high proportion of high-mannose-type *N*-glycans (HMGs) on their surface (20-22). These glycans are typically processed to hybrid and complex forms on glycoproteins secreted by healthy cells (23). Thus, the HMGs on the surface of HCV may be considered a druggable target. We have previously described the development of an HMG-targeting lectin-Fc fusion protein, or “lectibody”, called Avaren-Fc (AvFc), which was shown to bind with high affinity to clusters of HMGs on the HIV envelope protein gp120 and effectively neutralize multiple HIV clades and groups including HIV-2 and simian immunodeficiency virus (24). Further analysis indicated that AvFc can bind to HCV E2 protein (24). Therefore, in this study, we aim to investigate the anti-HCV therapeutic potential of AvFc in *in vitro* neutralization assays and an *in vivo* HCV challenge study using PXB-mice®, a chimeric uPA/SCID mouse model transplanted with human hepatocytes (25).

## MATERIALS AND METHODS

### Animal Care

The use of animals was approved by the University of Louisville**’**s Institutional Animal Care and Use Committee and the Animal Ethics Committee of PhoenixBio Company, Ltd. (Resolution No.: 2281). All animals were given a standard diet and water *ad libitum* and housed in a temperature and humidity-controlled facility with a 12-hour day/night cycle.

### Production of AvFc and non-HMG-binding AvFc variant

AvFc and a non-HMG-binding variant (AvFc^lec-^) were produced by agroinfiltration with magnICON® vectors in *Nicotiana benthamiana* plants as previously described (24). AvFc was purified from plants after a 7-day incubation period using protein A and ceramic hydroxyapatite (CHT) chromatography.

### HCV neutralization assays

Huh-7 cells (26) and HEK-293T cells (ATCC) were cultured in Dulbecco’s modified Eagle’s medium (DMEM) supplemented with 10% heat-inactivated fetal calf serum and 1% penicillin/streptomycin. To produce cell cultured HCV (HCVcc), we used a modified version of the plasmid encoding JFH1 genome (genotype 2a), provided by T. Wakita (National Institute of Infectious Diseases, Tokyo, Japan) (27, 28). The H77/JFH1 chimera, which expresses the core-NS2 segment of the genotype 1a polyprotein within a genotype 2a background, has been described previously (29). The genotype 4a ED43/JFH1 (30), genotype 5a SA13/JFH1 (31), and genotype 6a HK6a/JFH1 (32) infectious HCV recombinants were provided by J Bukh (University of Copenhagen, Denmark). Retroviral pseudotypes bearing HCV envelope glycoproteins of JFH1 virus (HCVpp) expressing the *Firefly* luciferase reporter gene were produced in HEK-293T as previously described (33). Inhibitory effects were determined by quantifying infectivity by indirect immunofluorescence with the anti-E1 mAb A4 (34) or an anti-NS5A polyclonal antibody kindly provided by M Harris (University of Leeds, UK).

### Formulation buffer optimization

Initial buffer screening was performed in 30 mM glutamate, acetate, citrate, succinate, histidine and phosphate buffers at pH 4.5 – 7.5 (see Supplementary Table 1). All the buffer agents were purchased from MilliporeSigma (Burlington, MA, USA). AvFc was diafiltrated and adjusted to 1 mg/mL (or 62.5 μM) in respective buffers. Stability was evaluated by SDS-PAGE following incubation for 2 weeks at 37°C. The melting temperatures of AvFc were determined by differential scanning fluorimetry performed on an Applied Biosystems StepOnePlus RT-PCR system as described previously (24). Briefly, AvFc formulated in various buffers at a concentration of 50 μM was mixed with a final concentration of 50x SYPRO® Orange (ThermoFisher Scientific, Waltham, MA, USA) in a 96 well template (USA scientific, Ocala, FL, USA). The melting temperature (*T*_m_) was determined by the vertex of the first derivative of the relative fluorescence unit values in the melt curves. AvFc formulated into the optimized histidine buffer or PBS was then concentrated to 10 mg/mL and incubated at 4°C or room temperature. Absorbance at 280 nm and 600 nm was measured immediately after concentration and then again after 16 and 72 h. A_280_ was measured after centrifugation of precipitate.

**Table 1:** IC_50_ values for AvFc and Avaren against HCVcc

| Virus | Genotype | Avaren IC<sub>50</sub> (nM) | AvFc IC<sub>50</sub> (nM) |
| --- | --- | --- | --- |
| JFH1/H77 | 1a | 529.28 ± 158.78 | 1.69 ± 0.39 |
| JFH1 | 2a | 484.62 ± 109.16 | 1.69 ± 0.78 |
| JFH1/ED43 | 4a | 204.27 ± 1.65 | 2.85 ± 0.91 |
| JFH1/SA13 | 5a | 148.86 ± 2.48 | 2.33 ± 0.13 |
| JFH1/HK6a | 6a | 114.95 ± 52.93 | 1.95 ± 0.78 |
| **Average:** |  | **269.39 ± 65.00** | **2.10 ± 0.60** |

### Pharmacokinetic analysis

A pharmacokinetic profile for AvFc was generated following a single 25 mg/kg i.p. injection in C57bl/6 mice (The Jackson Laboratory, BarHarbor, ME, USA; 8-week-old males and females; n=4 per time point) and sampling blood at 0.5, 1, 2, 4, 8, 12, 24 and 48 h post injection. The concentration of AvFc was then measured using an HIV gp120-coated ELISA. Briefly, a recombinant gp120 (HIV CM235, NIH AIDS Reagent Program) was coated overnight at 0.3 μg/mL followed by blocking with 5% dry milk-PBST. Serum samples at varying dilutions were incubated for 2 h followed by detection by a goat anti-human Fc-HRP secondary antibody (ThermoFisher Scientific). The plasma concentration of AvFc was calculated by interpolating from a standard curve. PK parameters were calculated using the PKSolver Microsoft Excel Add-on (35).

### Toxicological analysis and HCV challenge study in PXB-mice

The mouse model of toxicological analysis and HCV infection and toxicological analysis was performed in PXB-mice® (cDNA-uPA^wild/+^/SCID, cDNA-uPA^wild/+^: B6;129SvEv-Plau, SCID: C.B-17/Icr-*<u>scid/scid</u>* Jcl). These mice contain transplanted human hepatocytes with a replacement index of greater than 70% as determined by blood human albumin (h-Alb) measurements prior to virus inoculation (25). Mice were separated into 3 treatment groups: AvFc^lec-^ (25 mg/kg, n=5) for 11 doses, or AvFc (25 mg/kg, n=5 each) for 8 or 11 doses. Treatments were co-administered i.p. with virus inoculation (5 × 10^5^ copies/kg) on day 0 with a genotype 1a strain (PBC002) and every other day thereafter. The general conditions and body weights of the animals were monitored every other day, while serum HCV RNA and blood h-Alb were measured every 7 days by RT-PCR and latex agglutination immunonephelometry (LZ Test “Eiken” U-ALB, Eiken Chemical Co., Ltd.) respectively. Serum alanine aminotransferase 1 (ALT) levels were determined either using a Fujifilm DRI-CHEM NX500sV clinical chemistry instrument or by ELISA (Institute of Immunology Co., Ltd., Tokyo, Japan). At the study termination on day 35, animals were euthanized and subject to gross necropsy and general health. Blood was also drawn via cardiac puncture and used for ALT, HCV RNA and h-Alb analyses.

### Histopathological analysis of liver tissues

Hematoxylin and eosin-stained liver sections from 3-4 mice per group were generated by Nara Pathology Research Institute Co., Ltd. (Nara, Japan) and evaluated by pathologists at SkyPatho, LLC. All slides were examined by a blinded, board-certified veterinary pathologist under a light microscope (BX43, Olympus Corporation, Tokyo, Japan). The tissues were assigned a severity score for a number of characteristics based on the 5-point scoring system of the CDISC SEND Controlled Terminology where 0: unremarkable, 1: minimal, 2: mild, 3: moderate, 4: marked; 5: severe; and P: present.

### Statistical Analyses

Statistical significance was analyzed by the GraphPad Prism 6 software (La Jolla, CA, USA). Mouse body weights, Alb, ALT and HCV RNA levels were compared using a repeated measures two-way analysis of variance (ANOVA) with the Geisser-Greenhouse correction. Multiple comparisons between groups at each time point were conducted and corrected using the Tukey method with the threshold of significance set at p = 0.05. Liver:body weight ratios were compared using one-way ANOVA.

## RESULTS

### AvFc exhibits broad anti-HCV activity *in vitro*

Building on our previous observation that AvFc has affinity to a recombinant HCV E2 envelope protein (24), we first examined whether AvFc inhibits HCV infection *in vitro* using multiple genotypes of cell culture-produced virus (HCVcc) or pseudotyped virus (HCVpp). AvFc significantly blocked the infection of the human liver cell line Huh-7 by HCVcc from genotypes 1a, 2a, 4a, 5a, and 6a, with 50% inhibitory concentration (IC_50_) values in the low nanomolar range (**Table 1** and **Figure 1A**). Compared to Avaren monomer, AvFc overall showed approximately 2-log higher activity, while no inhibitory effect was observed for a plant-produced anti-HIV broadly neutralizing antibody VRC01 that shares the same human IgG1 Fc region with AvFc (36). Additionally, Avaren and AvFc, but not VRC01, effectively neutralized HCVpp harboring a murine leukemia virus backbone, suggesting that the lectin and the lectibody act as an entry inhibitor (**Figure 1B**).

**Figure 1:**
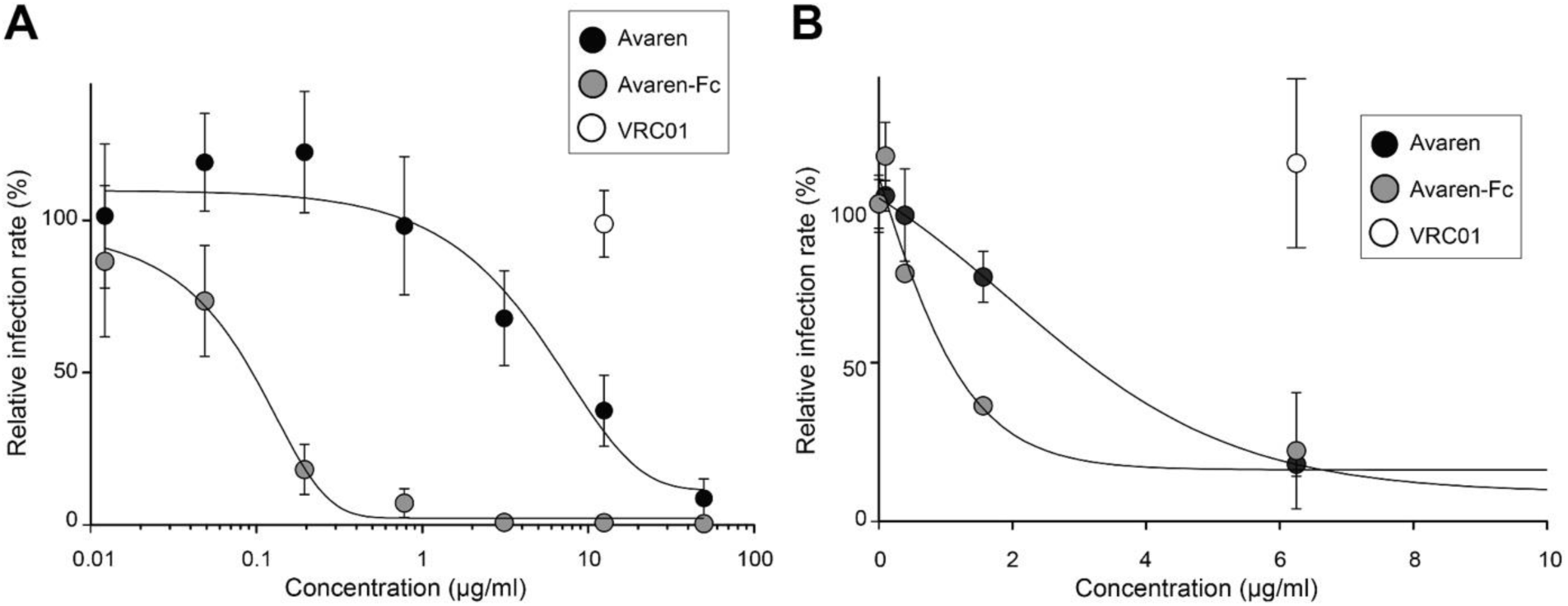
*In vitro* HCV inhibition assays. (A) Avaren and Avaren-Fc (AvFc) inhibit cell culture derived HCV. JFH1 virus was preincubated with Avaren, AvFc or the control antibody VRC01 for 30 min at 37°C before incubation with Huh-7 cells. At 48 h post-infection, infected cells were quantified by indirect immunofluorescence with an HCV-specific antibody. Results are expressed as percentage of infection compared to a control infection in the absence of compound. Error bars indicate standard errors of the mean (SEM) values from at least three independent experiments. (B) Avaren and AvFc inhibit HCV entry. Retroviral pseudotypes bearing HCV envelope glycoproteins of JFH1 virus (HCVpp) were preincubated with Avaren, AvFc or the control antibody VRC01 for 30 min at 37°C before incubation with Huh-7 cells. At 48 h post-infection, cells were lysed to quantify the luciferase activity. Results are expressed as percentage of infection compared to the control infection in the absence of compound. Error bars indicate SEM values from at least three independent experiments.

### Formulation of AvFc into a biocompatible buffer for *in vivo* studies

Previously, we found that AvFc has limited solubility in phosphate-buffered saline (PBS) at concentrations above 1 mg/mL (unpublished observation). In order to facilitate *in vivo* studies we screened for an optimal liquid formulation for systemic administration that can impart improved stability and solubility to AvFc at higher concentrations. Initial buffer screening revealed that AvFc is prone to degradation at and below pH 6.5, suggesting that AvFc is not stable in acidic pH conditions (**Figure S1**). Further pre-formulation studies led us to identify an optimal buffer composed of 30 mM histidine, pH 7.0, 100 mM sucrose and 100 mM NaCl. Although AvFc showed comparable *T*_m_ in the histidine buffer and PBS in differential scanning fluorimetry (62.49 ± 0.13°C vs. 62.68 ± 0.25°C; **Figure 2A**), SDS-PAGE analysis revealed that the lectibody holds superior stability in the histidine buffer upon accelerated stability testing via overnight incubation at 55°C (**Figure 2B**). When concentrated to ∼10 mg/mL, AvFc remained stable in solution in the histidine buffer over 72 h at 4°C and room temperature, while showing a significant concentration decrease concomitant with increasing turbidity in PBS (**Figure 2C**), further demonstrating the histidine buffer’s superiority for AvFc formulation.

**Figure 2:**
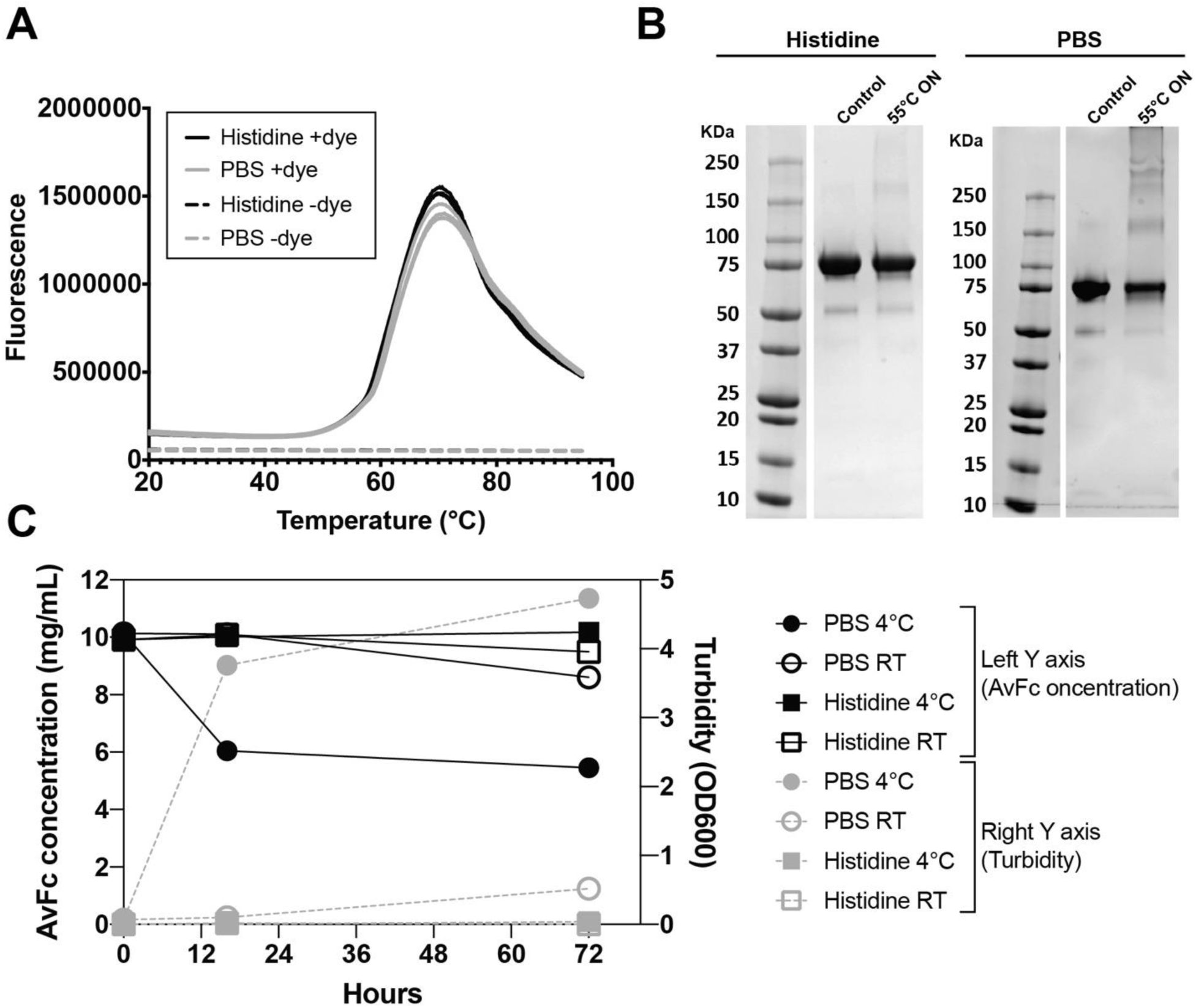
Liquid formulation development for AvFc. (A) Differential Scanning Fluorimetry (DSF) for melting temperature (*T*_m_) measurement. AvFc was prepared in 30 mM histidine buffer, 100 mM NaCl, 100 mM sucrose (“Histidine”; black line) or phosphate buffered saline (PBS; grey line) at a concentration of 1 mg/mL and analyzed in triplicate in the presence (solid line) or absence (dashed line) of the fluorescent dye SYPRO® Orange. *T*_m_ values were 62.49 ± 0.13°C in the Histidine buffer and 62.68 ± 0.25°C in PBS, as determined by the vertex of the first derivative of relative fluorescence unit values. (B) Accelerated stability testing of AvFc in the histidine buffer and PBS. AvFc, prepared at 1 mg/mL in the histidine buffer (“Histidine”; see above) or PBS were incubated overnight at 55°C, and 10 µg of the protein from each formulation was analyzed by SDS-PAGE under non-reducing conditions. A representative Coomassie-stained gel image is shown. The band at around 75 kDa corresponds to AvFc. Note that, after overnight incubation, PBS shows less band intensity for AvFc and more large-size aggregate bands than the histidine buffer. (C) Time course of concentration change and turbidity of AvFc solution in the histidine buffer and PBS. AvFc was formulated at 10 mg/mL in respective buffers and incubated at 4°C or room temperature (RT). After 16 and 72 h, the concentration was measured using a theoretical extinction coefficient at 280 nm of 1.6493 (mg/mL)^-1^ cm^-1^, whereas turbidity was assessed by absorbance at 600 nm. Representative data are shown for samples analyzed in triplicate.

### Pharmacological and toxicological analysis of AvFc in mice

To determine an optimal dosing regimen for an HCV challenge experiment, a pharmacokinetic analysis of AvFc was conducted in C57bl/6 mice. After a single i.p. injection of AvFc at a dose of 25 mg/kg, peak drug concentration was observed between 2 and 4 h, with a half-life of 24.5 h in male and 18.5 in female animals (**Figure 3**). After 48 h, in both male and female animals the plasma concentration of AvFc remained above a target trough concentration of 130 nM (10 μg/mL), at which AvFc showed >90% neutralization effects against HCV (see **Figure 1**). Consequently, these results suggested that administration of the drug every other day (Q2D) might be sufficient to keep the virus under control in a murine HCV challenge model.

**Figure 3:**
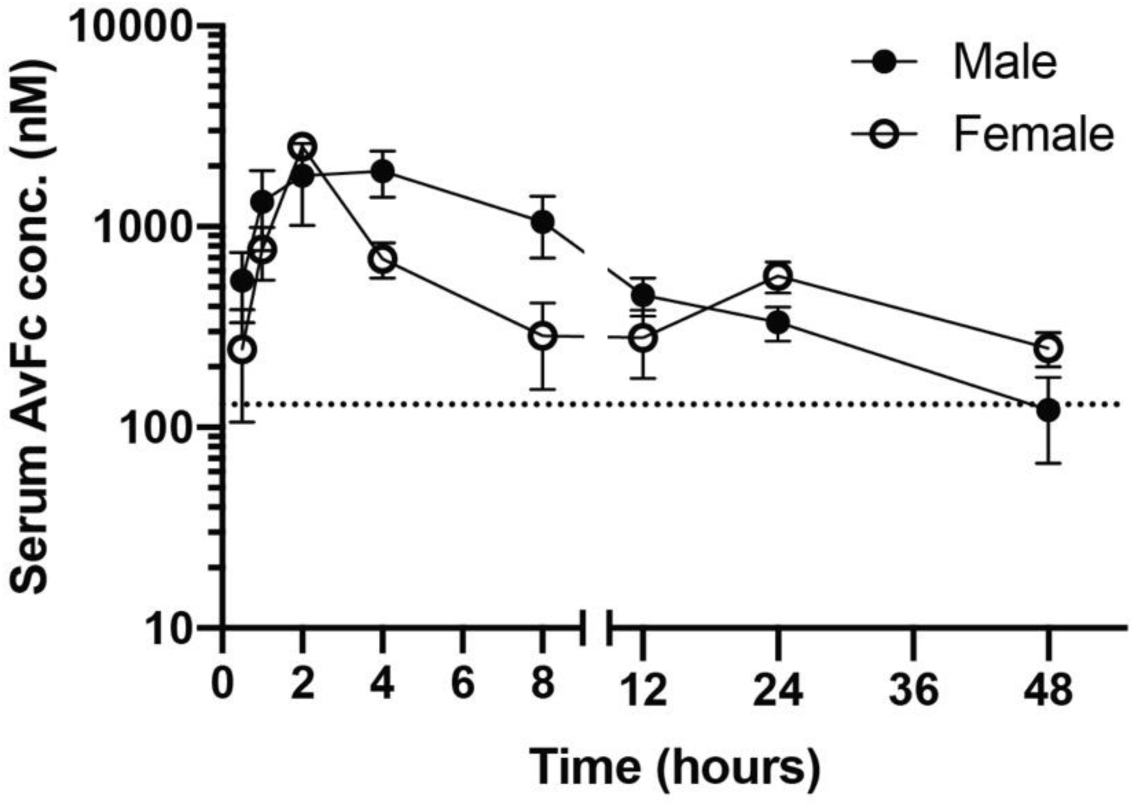
Pharmacokinetics of AvFc in Mice. AvFc pharmacokinetics were evaluated in C57bl/6 mice following a single i.p. injection of 25 mg/kg with blood sampled at various time points. Data are expressed as mean ± SEM from 4 mice per group. The average half-life was 24.5 h and 18.5h in male and female mice, respectively, as determined by the PKSolver Microsoft Excel Add-on. The peak concentration occurred between 2 and 4 h post administration. The target trough concentration of 130 nM (corresponding to 10 μg/mL) is indicated by a dashed line.

We then assessed the safety of Q2D administration of AvFc in PXB-mice®. To effectively discern potential toxicity associated with AvFc’s HMG-binding activity, we included an AvFc variant lacking HMG-binding activity as a control (AvFc^lec-^; **Figure S2**). PXB mice received either the vehicle (the histidine buffer described above) Q2D for 11 total doses, AvFc at 25 mg/kg Q2D for a total of 8 or 11 doses, or AvFc^lec-^ at 25 mg/kg Q2D for 11 total doses. As shown in **Figure 4A-C**, no significant differences in either body weights, blood h-Alb levels or serum ALT activity were observed. Additionally, no significant differences in relative liver weight were seen (**Figure 4D**). These results indicate that AvFc, formulated in the histidine buffer, is well tolerated in the immunocompromised mice engrafted with human hepatocytes.

**Figure 4:**
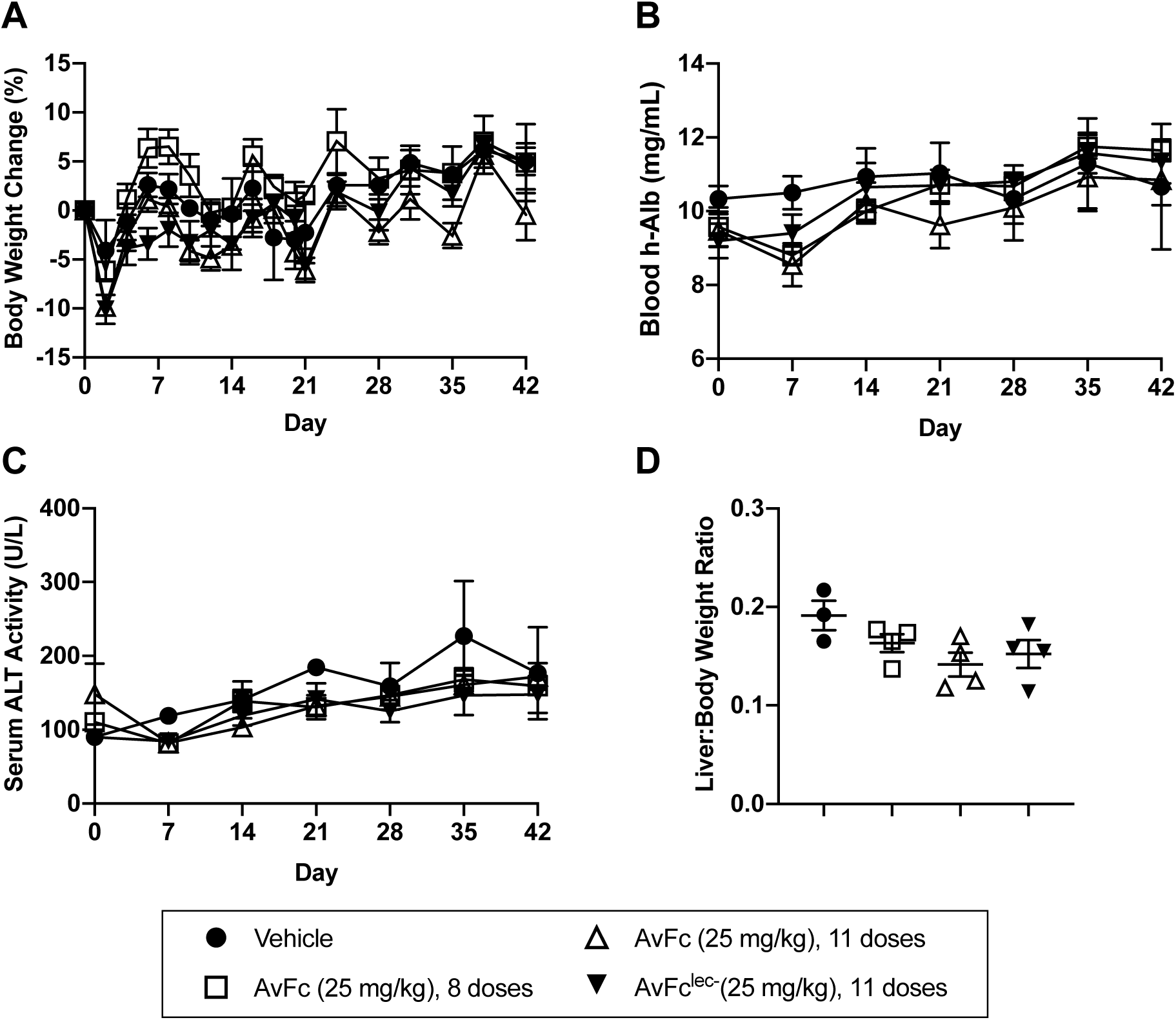
Toxicological analysis of systemically administered AvFc in the PXB® human liver chimeric mouse model. PXB mice were administered i.p. with AvFc or AvFc^lec-^ at 25 mg/kg (n=4 each), or the histidine buffer vehicle control (n=3) every 2 days (Q2D) and monitored for body weights, blood human albumin (h-Alb) levels and serum alanine aminotransferase (ALT) levels over 42 days. (A) Percent change of body weights from the initial day of dosing (Day 0). (B) Blood h-Alb levels. (C) Serum ALT levels. (D) Ratio of the liver weight to the body weight of individual mice at necropsy. Each data point represents mean ± SEM (A-C) and individual data with mean ± SEM (D) in each group. No significant changes in any of the safety endpoints were noted between the groups (A-C: two-way analysis of variance (ANOVA); D: one-way ANOVA).

Histopathology was performed to evaluate any potential toxicity to the human liver grafts due to AvFc administration (**Table 2** and **Figure 5**). In the human hepatocyte area, slight to moderate (score 2 to 3 in **Table 2**) macrovesicular fatty change, a characteristic change of human hepatocytes in the PXB-mouse, was observed in all mice including the vehicle-treated group (**Figure 5A-C**). Minimal inflammatory cell infiltration around vacuolated hepatocytes (Score 1) was seen in one mouse each from the 11 dose AvFc and AvFc^lec-^ groups (**Figure 5D, E**); however, this was unlikely treatment-related as a similar change is occasionally seen in PXB-Mice (PhoenixBio, unpublished observation). No AvFc treatment-specific change was observed, except for an incidental pigmentation in the Glisson’s sheath in one mouse (**Figure 5F**). Collectively, it was concluded that there was no treatment-related adverse effect in the liver tissue. The full pathology report may be found in the Supplementary Information.

**Table 2:** Histopathology of chimeric mouse liver tissue

|  | Vehicle |  |  | AvFc<sup>lec-</sup> |  |  |  | AvFc, 11 doses |  |  |  | AvFc, 8 doses |  |  |  |
| --- | --- | --- | --- | --- | --- | --- | --- | --- | --- | --- | --- | --- | --- | --- | --- |
|  | 101 | 102 | 103 | 201 | 202 | 203 | 204 | 301 | 302 | 303 | 304 | 401 | 402 | 403 | 404 |
| Mouse hepatocytes | 0 | 0 | 0 | 0 | 0 | 0 | 0 | 0 | 0 | 0 | 0 | 0 | 0 | 0 | 0 |
| Human hepatocytes |  |  |  |  |  |  |  |  |  |  |  |  |  |  |  |
| Fatty change, macrovesicular | 2 | 3 | 3 | 3 | 3 | 3 | 3 | 3 | 3 | 3 | 3 | 3 | 3 | 3 | 3 |
| Infiltrate, inflammatory cell, around vacuolated hepatocyte | 0 | 0 | 0 | 0 | 1 | 0 | 0 | 0 | 1 | 0 | 0 | 0 | 0 | 1 | 0 |
| Portal canal and others |  |  |  |  |  |  |  |  |  |  |  |  |  |  |  |
| Hepatocellular carcinoma, trabecular, with extramedullary hematopoiesis | P | 0 | 0 | 0 | 0 | 0 | 0 | 0 | 0 | 0 | 0 | 0 | 0 | 0 | 0 |
| Metaplasia, osseus | 0 | 2 | 0 | 0 | 0 | 0 | 0 | 0 | 0 | 0 | 0 | 0 | 0 | 0 | 0 |
| Pigmentation, brown, histiocyte, Glisson's sheath, focal | 0 | 0 | 0 | 0 | 0 | 0 | 0 | 0 | 0 | 0 | 0 | 1 | 0 | 0 | 0 |

**Figure 5:**
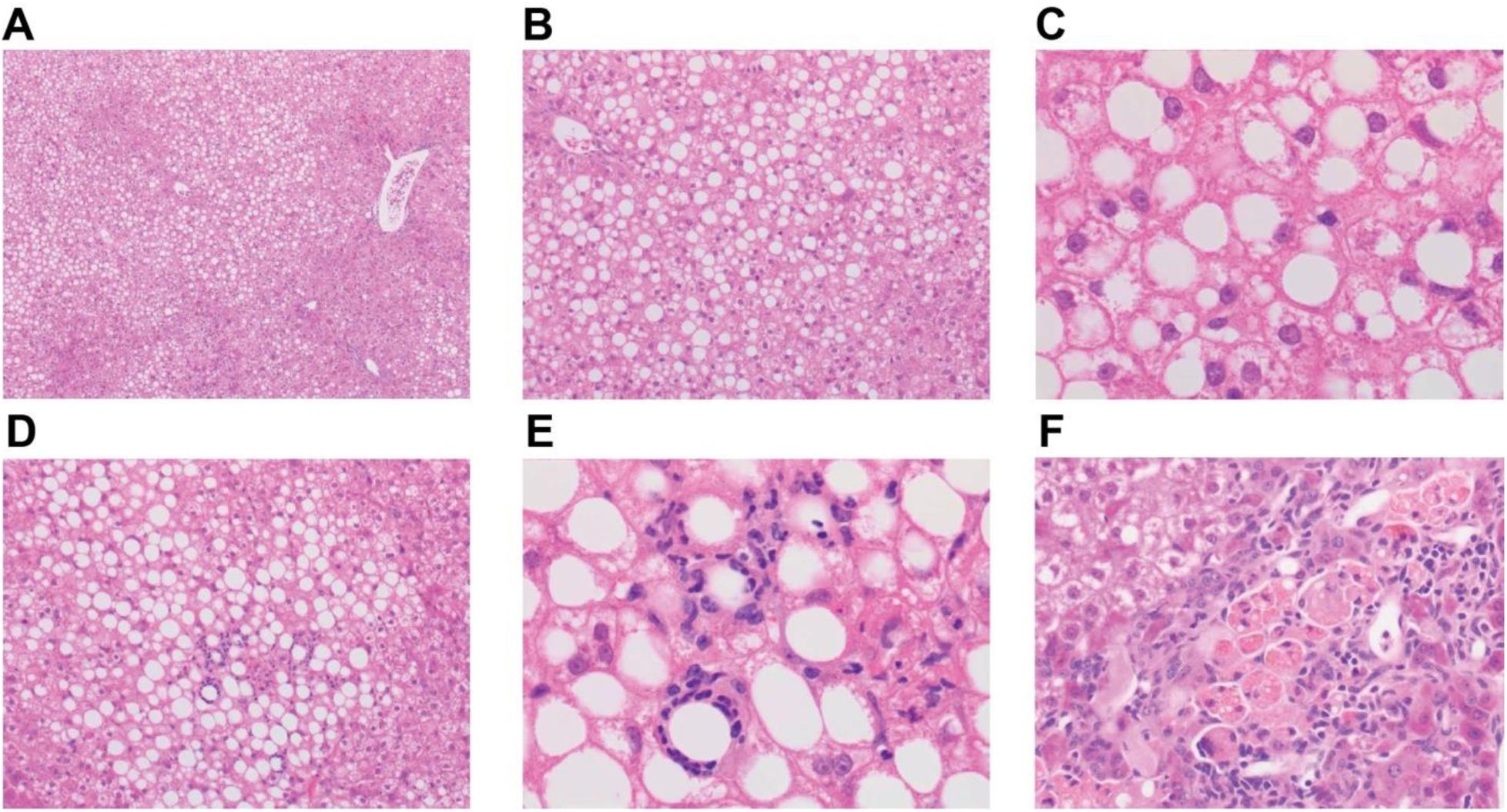
Histopathological examination of PXB mouse liver tissues. Representative hematoxylin/eosin-stained liver tissue section images corresponding to histopathological findings in Table 2 are shown. Liver tissues are from the toxicological study in Figure 4. (A) A 4x image from an animal in the vehicle control group (mouse ID: 103 in Table 2) showing low magnification of vacuolated hepatocytes. (B) A 10x image from a portion of panel A, containing many human hepatocytes with a large, well-defined rounded vacuole. (C) Higher magnification (40x) of panel B. (D) A 10x image from an animal in the AvFc^lec-^ group (ID: 202 in Table 2), showing small foci of inflammatory cell inflammation in the human hepatocyte area. (E) Higher magnification (40x) of panel D. Inflammatory cells appear to surround vacuolated hepatocytes. (F) A 20x image from an animal in the AvFc group (8 total doses; ID: 401 in Table 2). Histiocytic brown pigmentation in the Glisson’s sheath is noted only in this mouse.

### AvFc protects against HCV infection *in vivo*

Lastly, we assessed AvFc’s protective efficacy against HCV infection *in vivo* using the treatment regimen described above. PXB mice were inoculated i.p. with a genotype 1a virus along with initial treatment with 25 mg/kg of AvFc or AvFc^lec-^ on day 0. As shown in **Figure 6A**, AvFc^lec-^-treated mice showed high serum HCV RNA levels from day 7 post challenge through the end of the study on day 35. In sharp contrast, animals treated with both 8 and 11 doses of AvFc did not show any detectable level of HCV RNA in sera, indicating that the lectibody prevented the infection of human liver grafts by the virus. Similar to the results in Figure 3, overall no major toxicity signal was noted in body weights, h-Alb or h-Alt levels between the test groups although there was a temporal drop in body weight and h-Alb in one of the AvFc-treated group at an early timepoint, indicating that the liver grafts remained functional over the course of the study (**Figure 6B-D**).

**Figure 6:**
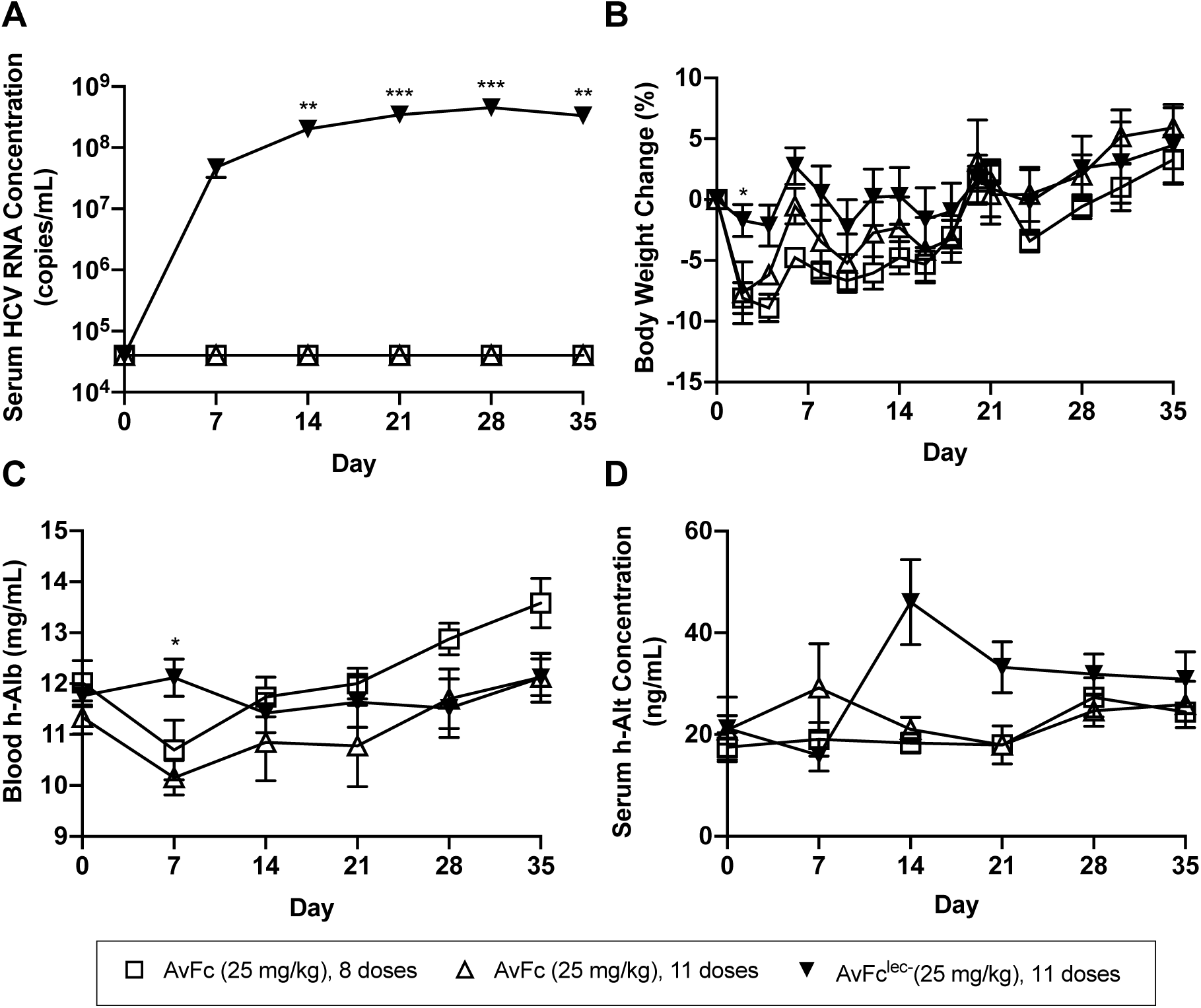
The protective effect of AvFc against HCV challenge in PXB mice. PXB mice were challenged i.p. with a HCV genotype 1a virus on Day 0 simultaneously with an initial treatment i.p. with either 25 mg/kg of AvFc or AvFc^lec-^. Treatment was continued Q2D for a total of 8 or 11 doses for AvFc and 11 doses for AvFc^lec-^ (n=5 each). The general conditions and body weights of the animals were monitored every other day, while serum HCV RNA and blood h-Alb were measured every 7 days. (A) Serum HCV RNA levels. AvFc treatment (both 8 and 11 doses) showed no detectable HCV RNA at any time point. **, ***p < 0.01, 0.001 (AvFc^lec-^ vs. both AvFc 8 and 11 doses); two-way ANOVA with Tukey’s multiple comparison test. (B-D) Time course of body weight change from day 0 (B), blood h-Alb levels (D) and serum h-Alt concentrations (D). Each data point represents mean ± SEM in each group. *p < 0.05 (AvFc^lec-^ vs. AvFc 8 doses in B and AvFc^lec-^ vs. AvFc 11 doses in C]; two-way ANOVA with Tukey’s multiple comparison test. No significant difference between groups at any timepoint was noted in D.

## DISCUSSION

In this study we demonstrated that the HMG-binding lectibody AvFc exhibits broad genotype-independent anti-HCV activity. Additionally, systemic administration of AvFc effectively protected chimeric human-mouse liver mice from infection with a genotype 1a virus without apparent toxicity, providing the first *in vivo* proof-of-concept for the lectibody’s antiviral potential.

The mechanism of HCV neutralization by AvFc is likely through binding to HMGs on the E1/E2 envelope protein dimer, which blocks their interaction with host cell receptors and viral entry. Unlike HIV envelope glycoproteins, whose glycan content can vary widely between strains, the number and position of glycosylation sites on E1/E2 are highly conserved, indicating their critical role in HCV’s infectious processes (37). The notion that AvFc functions as an entry inhibitor is supported by the facts that the lectibody has affinity to the E2 protein (24) and that other mannose-binding lectins, such as Griffithsin or Cyanovirin-N, inhibit entry in this manner (38, 39). AvFc inhibited multiple genotypes of HCV with an average IC_50_ over 100-fold lower than that of the monomer Avaren lectin (**Table 1**), indicating that the multivalent recognition of HMGs on the surface of the virus, brought about by the dimerization of Avaren via Fc fusion, led to greater entry inhibition. Unlike other antiviral lectins, however, the inclusion of the human IgG1 Fc region implicates the possibility of Fc-mediated effector functions, such as antibody-dependent cell-mediated cytotoxicity, against infected cells. In fact, Fc-mediated effector functions greatly contributed to the antiviral potency of AvFc against HIV, as determined by a primary cell-based inhibition assay and an antibody-dependent cell-mediated viral inhibition assay (24). Accordingly, the remarkable efficacy seen in the present *in vivo* HCV challenge study may be partially Fc-mediated. Further investigations are necessary to address this possibility.

The present study also demonstrated that AvFc therapy is well tolerated in mice and human hepatocytes, as Q2D i.p. administration of 25 mg/kg of AvFc up to 11 doses did not show any obvious toxicity in PXB mice by gross necropsy or histopathology of engrafted human hepatocytes, nor did it result in significant changes in body weight, h-Alb, or ALT levels (**Figure 3, 4**). We hypothesize that the lack of any significant toxicity is attributable to AvFc’s unique HMG-binding mechanism, whereby it requires multivalent interaction with several HMGs in proximity to exhibit high affinity binding to a glycoprotein target. In line with this hypothesis, Hoque et al. demonstrated that the three binding pockets of the parent lectin actinohivin can bind up to three independent HMGs, providing high affinity binding when the HMGs are in relatively close proximity (40). This implies that AvFc may not effectively interact with healthy normal cells and tissues that do not usually exhibit clusters of HMGs on their surfaces. In contrast, glycoproteins of many enveloped viruses display a high proportion of these immature forms of *N*-glycans (20-22). While HCV E2 has fewer *N*-glycosylation sites (around 11) than the HIV glycoprotein gp120 (which has between 20 and 30 depending on the strain), E2 is likely present on the surface of HCV at a higher density and thus provides higher local concentrations of HMGs (41). Further studies are necessary to reveal a threshold HMG concentration which enables efficient interaction between AvFc and the surfaces of cells or viruses.

While alcoholic liver disease has now surpassed HCV infection as the number one indication for liver transplantation in the US, a large number of procedures will continue to be performed for the foreseeable future in patients with HCV-related decompensated cirrhosis (42). A major outstanding issue is the lack of effective treatment protecting the allograft liver from recurrent infection by the virus that remained circulating in the periphery at the time of transplant. As a consequence, reinfection of donor livers universally occurs, as early as in the first 90 minutes upon reperfusion (17), and can result in accelerated fibrosis and increased risk of graft failure, cirrhosis, and hepatocellular carcinoma (43). In fact, allograft failure due to reinfection is the leading cause of secondary transplants and death in HCV-infected patients who have received liver transplant (44), Patients cured of HCV with DAAs after liver transplantation still have a higher than normal risk of hepatocellular carcinoma (45), and the high cost of the drugs represents a significant barrier to their widespread use. Furthermore, emergent drug resistance even in DAA combination therapies, though rare, represents a particular challenge for further treatment (46). Thus, while the effectiveness of DAAs is not in question, there are still unmet needs that may be addressed through the use of entry inhibitors. As of yet, no entry inhibitor has been approved for the treatment or prevention of HCV. Two major drug candidates, Civacir® and MBL-HCV1, have shown some promise in clinical trials (NCT01804829, NCT01532908) (47, 48). Though larger studies are needed, it appears that entry inhibitors in combination with DAAs may represent a new treatment paradigm for HCV patients receiving liver transplant. Despite that both MBL-HCV1 and Civacir® are capable of neutralizing a broad range of HCV genotypes, viral resistance can still develop through mutations in the envelope proteins E1/E2, in particular through shifting glycan positions (49, 50). In this regard, AvFc in its own right could be less susceptible to amino acid mutations because it targets the glycan shield of the virus rather than a specific epitope. Deletions of glycans, even if occurring following prolonged exposure to a carbohydrate-binding agent like AvFc, may result in significant decrease in viral fitness by decreasing E1/E2 incorporation into HCV particles or increased susceptibility to humoral immunity due to breach in the glycan shield (37, 51). Our results provide a foundation to test the above hypotheses and feasibility of the HMG-targeting anti-HCV strategy.

In conclusion, the present study provided an important proof of concept for the therapeutic potential of AvFc against HCV infection via targeting envelope HMGs. In particular, the lectibody may provide a safe and efficacious means to prevent recurrent infection upon liver transplantation in HCV-related end-stage liver disease patients.

## ACKNOWLEDGMENTS

We thank Adeline Danneels, Lucie Fénéant, Czeslaw Wychowski, Lauren Moore and Jessica Jurkiewicz for their experimental help. We are also grateful to R. Bartenschlager, J. Bukh, F.L. Cosset, M. Harris and T. Wakita for providing essential reagents. The immunofluorescence analyses were performed with the help of the imaging core facility of the BioImaging Center Lille Nord-de-France.

## FUNDING

This work was supported by the University of Louisville ExCITE program, which was funded by U.S. National Institutes of Health (U01 HL127518) and the Leona M. and Harry B. Helmsley Charitable Trust. AvFc^lec-^ was created from work supported by a National Institutes of Health grant (R33 AI088585).

### Transparency declarations

N.M. is an inventor on a patent concerning the composition and utility of AvFc (U.S. patent number 8802822).

